# Detecting sex-linked genes using genotyped individuals sampled in natural populations

**DOI:** 10.1101/2020.01.31.928291

**Authors:** Jos Käfer, Nicolas Lartillot, Gabriel A.B. Marais, Franck Picard

## Abstract

We propose a method, SDpop, able to infer sex-linkage caused by recombination suppression typical of sex chromosomes. The method is based on the modeling of the allele and genotype frequencies of individuals of known sex in natural populations. It is implemented in a hierarchical probabilistic framework, accounting for different sources of error. It allows to statistically test for the presence or absence of sex chromosomes, and to infer sex-linked genes based on the posterior probabilities in the model. Furthermore, for gametologous sequences, the haplotype and level of nucleotide polymorphism of each copy can be inferred, as well as the divergence between both. We test the method using simulated and human sequencing data, and show that, for most cases, robust predictions are obtained with 5 to 10 individuals per sex.

## 1 Introduction

Sex chromosomes, which are found in many species with genetic sex determination, are key elements of the biology of the sexes (e.g. between-sexes differences in development, morpho-anatomy, physiology) and are consequently studied by different branches of biology (e.g. genetics, genomics, developmental biology, physiology, evolutionary biology, medical research, agronomy). While sex chromosomes share striking features even between independently evolved systems, there is also much diversity (reviewed in Bachtrog et al., 2014).

Sex chromosomes are thought to evolve from a pair of autosomes (Lahn and Page, 1999; Charlesworth et al., 2005). One sex is characterized by a pair of sex chromosomes for which recombination is suppressed over part of the length, while the other has two identical chromosomes that recombine normally. Thus, the sex chromosome which is limited to one of the two sexes (the so-called “‘sex limited chromosome”‘, SLC) contains a part that never recombines, and evolves independently from the homologous part on the other sex chromosome. These systems are termed according to the sex that is heterogametic; when the males are the heterogametic sex, the sex chromosomes are named X and Y, and when the females are heterogametic, they are named Z and W; Y and W are the SLCs in these systems, respectively. The recombining part of the sex chromosomes is called the pseudo-autosomal region.

The complete lack of homologous recombination in a part of the SLC has dramatic consequences for its evolution (Charlesworth et al., 2005; Lynch, 2007): it accumulates deleterious mutation (Muller’s ratchet) and transposable elements, which deteriorate gene function and finally chromosome integrity. Once most genes have lost their function, parts of the chromosome can be lost without negative effects on the individual’s fitness, leading to situation that is considered paradigmatic of a large X chromosome and a small Y chromosome with only few functional genes, as in humans. However, in many sex chromosome systems, the SLC does not differ significantly in size from its homologous counterpart, and recombination is suppressed in only a small region of these chromosomes. As more sex chromosome systems are being studied in phylogenetically distant lineages, more questions arise on their similarities and differences (e.g. Charlesworth, 2016; Muyle et al., 2017; Ponnikas et al., 2018).

Even though sequencing technologies have been improving constantly, sex chromosomes have remained difficult to study for a long time. In particular, SLC sequences used to be scarce for two reasons (Tomaszkiewicz et al., 2017). First, genome projects have been typically using whole-genome shotgun (WGS), in which DNA is fragmented and sequenced using short read technologies and then re-assembled in contigs. The assembly of the non-recombining regions of the SLC, which are full of repeats, using short reads is virtually impossible (Hughes and Rozen, 2012). Second, in the heterogametic sex, read coverage for the sex chromosomes is half the coverage for the autosomes, making the assembly of both sex chromosomes more difficult, while sequencing of the homogametic sex only yields one of the chromosomes.

Recently, the development of new methods to study sex chromosomes has boosted their study. The first Y sequence of humans and other species (e.g. papaya) have been obtained with a specific methods based on BACs, such as the SHIMS, which are very time-consuming and costly (Skaletsky et al., 2003; Hughes and Rozen, 2012; Wang et al., 2012; Bellott et al., 2014; for example, the human Y sequencing costed 6 million dollars). Costs have decreased in a new version of SHIMS, using Illumina short-read sequencing of pooled BACs instead of Sanger sequencing of individual BACs (Bellott et al., 2018), but they are still substantial for any Y chromosome larger than a few megabases. More efficient and cheaper methods have thus been developed (reviewed in Muyle et al., 2017; Palmer et al., 2019).

A first alternative approach was to compare male and a female genomes obtained using short reads (WGS). Absence in the female genome is enough to identify the Y contigs (Carvalho and Clark, 2013; Cortez et al., 2014). The X contigs can be identified by computing the read depth in male and female: the male to female read depth ratio is expected to be 1 for autosomes and 0.5 for the X, as males have one X copy and females two (Vicoso and Bachtrog, 2011). Variants of this approach have been widely used (e.g. Vicoso et al., 2013a,b; Hall et al., 2013; Li et al., 2018). However, this approach works only with strongly diverged, old sex chromosomes, in which the reads map uniquely to the X or the Y chromosomes but not to both. In more recently evolved, weakly diverged systems, the X and the Y still share a high level of sequence similarity, and reads cannot be unambiguously assigned to one of the sex chromosomes with alignment methods.

To study weakly divergent sex chromosomes, a second category of empirical methods uses segregation patterns in genotyping data from natural populations. In XY systems, an excess of heterozygotes is expected in XY males compared to XX females at the sex-linked genes. Such patterns could be identified using male versus female *F*_*ST*_ (e.g. Zhou et al., 2017) or differences in heterozygote frequencies (e.g. Picq et al., 2014; Kirkpatrick and Guerrero, 2014). GWAS using sex as the studied trait is another option which will work based on different allele frequencies in the sexes, but a precise model of the expected association is preferable (Galichon et al., 2012). Indeed, the patterns these empirical approaches look for can be caused by other processes than sex linkage, such as sex-antagonistic selection (e.g. Qiu et al., 2013). Such methods thus have high false positive rates and either must use conservative cut-offs, which significantly reduces their sensibility, or are applicable to species with well-assembled genomes (e.g. Picq et al., 2014; Mathew et al., 2014). An empirical approach somewhat intermediate between the former and the latter categories is to identify sex-specific sequences in the genomic reads, but these mostly only allow a first characterization of the sex chromosome system, and further sequencing is needed to study the sex-linked regions (Scharmann et al., 2017; Torres et al., 2018).

Recently, the SEX-DETector statistical framework, relying on the modeling of the transmission of alleles that account for the observed genotypes, was introduced (Muyle et al., 2016). The method has been shown to yield high power and high sensitivity with as few as five offspring of each sex, and has successfully been used on a number of species (Muyle et al., 2018; Veltsos et al., 2019; Martin et al., 2019; Prentout et al., 2020). It can be used for several purposes, using a single dataset: testing whether a species has sex chromosomes and of which kind (XY or ZW), identifying sex-linked genes, reconstructing the haplotypes of the gametolog copies, and estimating expression of each of the copies if RNAseq data are used. However, its requirement to produce a controlled cross for sequencing limits its use to species than can be easily bred or cultivated in controlled conditions, and hinders its application to species with long generation times.

A method that can be applied to sequencing data from individuals sampled in natural populations, DETSEX, was introduced by Gautier (2014). While it offers some attractive features, like automatic sexing of some individuals in the sample, it relies on ploidy levels to detect sex-linked sequences, and can thus only be used for sufficiently divergent sex chromosomes. The method has, to our knowledge, not been used in any study that aimed to identify sex-linked sequences.

We here introduce a new hierarchical framework based on the modeling of genotype and allele frequencies in a population, according to autosomal and sex-linked segregation types. As SEX-DETector statistically analyses Mendelian segregation and its deviations due to sex-linkage, the new method does so based on the Hardy-Weinberg equilibrium and its deviations. The method we present here is “‘SEX-DETector in a population”‘, and is thus termed “‘SDpop”‘. It can be applied to any sample collected from natural populations, provided the sex of the individuals can be determined morphologically. Despite the similarity of its goals with those of SEX-DETector (notably, applicability to sex chromosome systems of any age, statistical testing, likelihood-based inferences, prediction of gametolog haplotypes), it is an entirely different model due to the data it handles, the underlying population genetic expectations, and some of the predictions it can produce.

## 2 Materials and methods

### 2.1 Model

#### 2.1.1 Terminology

A sex-linked gene (i.e. a gene that is present on the non-recombining region of the sex chromosomes) has two independent copies. These copies will accumulate mutations, be subject to selection and drift, and accumulate allele fixations independently from one another. These copies thus diverge after the suppression of recombination, in a way that is reminiscent of the divergence of paralogous genes after gene duplication; the two copies of a sex-linked gene are thus termed “‘gametologs”’ (Garcia-Moreno and Mindell, 2000). Eventually, due to neo- or sub-functionalization, or the accumulation of deleterious mutations, one of the copies can get lost, most often the copy present on the Y or W chromosome. The remaining gene, on the X or Z, is termed “‘X-”’ or “’ Z-hemizygous”‘.

Gametology and hemizygosity cause differences in the relation between genotype and allele frequencies with respect to autosomal segregation. More precisely, these differences occur in one sex, i.e. the heterogametic one. If one considers a sexual, random mating population of diploid individuals, most autosomal genes will be at Hardy-Weinberg equilibrium. The deviations from this equilibrium in the heterogametic sex are themselves equilibria, that can be described assuming panmixia, and that are the basis of the present model. For gametologs, two independent copies are present in the heterogametic sex, while for hemizygous genes, only one copy is present. For non-sex-linked genes, the presence of two independent copies can also occur for both sexes under paralogy, and the presence of only one copy for the haploid mitochondrial and chloroplastic genes, or when only one allele is expressed in transcriptome data. We thus distinguish five possible segregation types in a diploid population: diploid autosomy (hereafter just called “’ autosomy”‘), haploidy, paralogy, hemizygosity, and gametology.

We model genotypes that are obtained by mapping sequencing data on a reference sequence. As the organisms are supposed to be diploid, the input data at all positions are diploid genotypes, and deviations from diploidy will be detected afterwards. We consider polymorphisms consisting of two alleles; these can be single nucleotide polymorphisms (SNPs) or structural variants (indels and length polymorphisms). For paralogs and gametologs, which are the result of the mapping of two non-recombining copies, we assume that cases in which both copies are polymorphic at the same position and for the same two alleles are very rare and can be neglected.

#### 2.1.2 Population genetic equilibria

We adopt a hierarchical probabilistic framework in which the distribution of alleles is modeled given a segregation type for each polymorphism. A technical presentation and derivations are given in the Appendix; here, we describe the general principles of the model.

Sex-linkage produces different genotype-allele equilibria for each sex. We consider sites with two alleles, *a* and *b*, and three possible genotypes *g*′ ∈ {*aa, ab, bb*}. The genotype frequencies are denoted *p* (*g* ′) if they are equal for both sexes, and indexed with ♂ and ♀ symbols when different. In the following, the allele frequency *f* is the frequency of allele *a*, unless otherwise stated.

For **autosomal segregation**, we can simply write

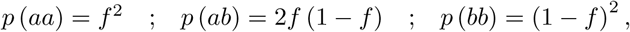

which is the Hardy-Weinberg equilibrium.

**Haploid genes** correspond to haploid genotypes, but genotyping will have given diploid genotypes. No sex-specific difference is expected, so

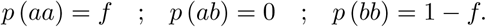

**Paralogy** is caused by the mapping of the reads of two more or less recently duplicated genes on one locus in the reference. Thus, paralogous genes have tetraploid genotypes, which again are considered diploid by the genotyper. There is no recombination between the copies, that thus evolve independently. Only biallelic sites are modeled here; for simplicity, we assume that one of the copies is fixed for one of the alleles. Thus, if one copy is fixed for allele *a*, the tetraploid genotypes are *aaaa, aaab* and *aabb*; the latter two will yield the diploid genotype *ab*. We define the allele frequency *f* as the frequency of allele *b* in the polymorphic copy. Again, no difference between the sexes is expected, and we thus obtain the diploid genotype frequencies

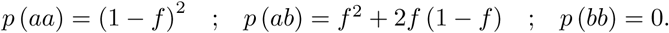

A choice has to be made about which allele to consider fixed, as this is not always clear *a priori*. The previous segregation types, autosomal and haploid, are symmetrical with respect to the choice of the allele to model, *i*.*e*., the expected genotype frequencies are strictly identical whether the frequency of allele *a* was used or the frequency of allele *b*. The paralogous segregation type is asymmetrical with respect to this choice: considering allele *a* or allele *b* as fixed does not lead to identical genotype frequencies. The details about the calculations are specified in the Appendix.

For **hemizygously segregating genes**, the members of one sex are haploid while the others are diploid. Thus, in one sex, the Hardy-Weinberg equilibrium is expected, while in the other, the expectations are the same as for the haploid segregation type. In the case of an XY system, expectations are thus

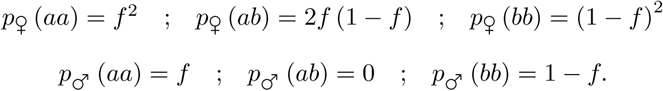

**Gametologous segregation** is characterized by the presence of two independent copies in one sex, and one copy in the other. As for the paralogous case, we assume that an allele is fixed in at least one of the copies. We have to distinguish two cases: either the X copy is polymorphic, or the Y copy. In either case, we have to choose which allele we consider fixed on one of the copies, thus leading to four different allele-genotype equilibria. Here, one case of X-polymorphism and one case of Y-polymorphism are described; details about the four equilibria are given in the Appendix.

In the case of polymorphism on the X, female genotypes are modeled by the Hardy-Weinberg equilibrium, while male genotypes show a specific deviation. If we consider allele *b* to be fixed on the Y chromosome, and define *f* as the frequency of *a* on the X chromosome, we get

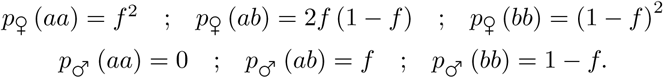

For a polymorphism on the Y chromosome, considering allele *b* to be fixed on the X-chromosome and defining *f* to be the frequency of allele *a* on the Y chromosome yields

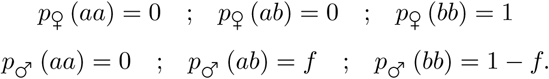

The X-hemizygous and XY segregation described here can easily be converted to Z-hemizygous and ZW segregation (see Appendix for details).

#### 2.1.3 Parameters and optimization

SDpop provides a complete probabilistic model for the posterior inference of segregation types at the level of polymorphisms. The input of SDpop consists in the observed genotype (*g*) frequencies at each biallelic site, that can differ from the (unobserved) true genotypes *g*′ with probability *e*. Then we introduce the segregation type, modeled as an unobserved variable *S* with multinomial *prior* distribution of parameter *π*. The conditional probabilities of the true genotypes under each segregation type, *p* (*g*′), are then obtained through the population genetic equations described above, using the empirical allele frequencies. This yields directly the conditional observed genotype probability *p* (*g* |*S*) = *p* (*g*′|*S*) *p* (*g*| *g*′). Our model also considers that under gametologous segregation, either one of the gametolog copies could be polymorphic, as well as the fact that in the gametologous and paralogous segregation types, either one of the alleles can be fixed in one of the copies, with equal prior probability (see Appendix). Thanks to an Expectation-Maximization (EM) algorithm we infer the parameters of the model by maximum likelihood, and provide the posterior probability of each segregation type at each polymorphic, *p*(*S*| *g*), site using Bayes’ rule. We also propose a contig-wise version of this posterior, by aggregating the site-wise posterior probabilities using geometric means of site-wise posteriors. All the technical derivations of SDpop are provided in the Appendix.

SDpop can investigate several models of sexual systems: absence of sex-linkage, sex-linkage of the XY type, sex-linkage of the ZW type, and sex-linkage of both types. Apart from the sex-linkage, models can be run with or without haploid and paralogous segregation types. The XY and ZW model include both hemizygous and gametologous segregation types. The likelihood of the model when parameter value convergence has been attained is used to calculate the Bayesian Information Criterion (BIC), which allows to compare models with different numbers of parameters.

#### 2.1.4 Haplotypes and population genetics inferences

For the sequences inferred as gametologous, several statistics can be calculated from the optimized values. The posterior probabilities of the segregation subtypes indicate which one of the copies (X or Y) is polymorphic, and which one of the alleles is fixed in one of the copies. We can thus infer the frequency of the alleles in the X and Y copies (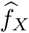 and 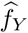), which is the mean of the empirical allele frequencies, weighted by the posterior probability for each subtype. These allow to reconstruct the X and Y haplotypes, by using only alleles that are fixed (or nearly so) in the X and Y copies.

Furthermore, the inferred allele frequencies allow to calculate the nucleotide diversity of the X and Y copies (*π*_*X*_ and *π*_*Y*_) and divergence between both gametolog copies (*D*_*XY*_) as the means of the diversity and the divergence over the whole contig:

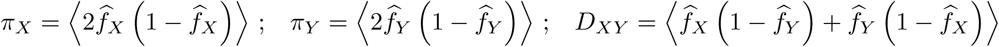

#### 2.1.5 Implementation and availability

SDpop is written in C with some C++ functionalities. It is available on https://gitlab.in2p3.fr/sex-det-family/sdpop.

### 2.2 Simulations

Sets of *K* contigs of *n* individuals per sex (i.e., 2*n* individuals in total) were generated using *ms* (Hudson, 2002), that simulates samples drawn from a population under the neutral Wright-Fisher model of genetic variation. Samples of autosomal contigs were generated by simulating 4*n* haploid sequences and combining them arbitrarily into 2*n* diploid samples. The parameter *θ* = 4*N*_*e*_*µ* = 4*N*_*e*_*uL* gives the average number of segregating sites. Here, *µ* is the overall mutation rate for the sequence with length *L* and per base mutation rate *u*. The heterozygosity rate (per site) is thus 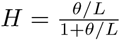, which is equal to the level of polymorphism *π*. Thus, for example, to simulate a sequence of *L* = 500 base pairs with a level of polymorphism *π* = 0.002, we use the parameter 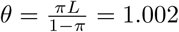.

Gametologous contigs were generated from a sample of 3*n* X-linked sequences and *n* Y-linked sequences, that were simulated by supposing two populations that split at time *t* from a population with size *N*_*e*_ into one population of size 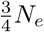 (X-linked sequences) and one of size 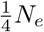 (Y-linked sequences). In `ms`, the time *t* is scaled by 4*N*_*e*_, such that, according to coalescent theory, *t* = 1 is the average time to fixation for a neutral mutation. X-hemizygous sequences were simulated similarly, except that no Y-linked sequences were simulated; male “‘diploid”’ genotypes were obtained from the haploid X sequence for each individual.

Errors were introduced after the coalescent simulations, such that each individual at each position had a probability *e* to have an erroneous genotype. Two types of errors were introduced: either homozygous genotypes were rendered heterozygous, or heterozygous genotypes homozygous, both with the same rate. As there are typically many more homozygous than heterozygous genotypes, many more errors occur on monomorphic sites than on polymorphic sites. SDpop estimates parameters using polymorphic sites only, and in this subset of all sites, the fraction of sites with errors is higher than in the whole genome, which includes monomorphic sites as well. Thus, the dataset of polymorphic sites on which SDpop runs is enriched in sites with errors; we refer to the error rate among polymorphic sites as the “‘effective error rate”‘, *e*_*e*_. As an example, for a series of simulations with 2, 5, 10, 20, 50, and 100 individuals per sex, which were all conducted with an error rate of 0.0001 and a level of polymorphism (*π*) of 0.002, *e*_*e*_ is 0.015, 0.011, 0.0088, 0.0068, 0.0045, and 0.0030, respectively.

The four models of SDpop (without sex chromosomes, with XY chromosomes, with ZW chromosomes, and with both XY and ZW chromosomes) were run on each simulation. The best model was chosen based on the BIC values. The contigs were classified as sex-linked or not based on SDpop’s posterior probabilities of the contigs, using a posterior probability threshold of 0.8, and this classification was used to count the number of true positives (TP), false positives (FP), and false negatives (FN). To test the power and precision of the classification of contigs as sex-linked or not is quantified by the true positive rate (TPR = TP*/* (TP + FN)) and the positive predictive value (PPV = TP*/* (TP + FP)). Nucleotide polymorphism and divergence were calculated using the observed allele frequencies before errors were added.

### 2.3 Human data

Exome-targeted sequence data for Finnish individuals (Auton et al., 2015) were downloaded from the 1000 genomes project website. To facilitate file handling in the analysis pipeline, only individuals for which all reads were present in two fastq files (forward and reverse) were retained, i.e. a total of 66 individuals (44 females and 22 males). These reads were mapped on the human genome reference GRCh37 after removal of the Y sequence was removed (the reference used includes the primary assembly, i.e. chromosomal plus unlocalized and unplaced contigs, and the rCRS mitochondrial sequence, Human herpesvirus 4 type 1 and the concatenated decoy sequences) using BWA (Li and Durbin, 2009) with standard parameters. The individual alignments to the reference were merged, and the genotypes were called using bcftools (mpileup & call), and the targeted exons (file provided by the 1000 genome project) were extracted using bedtools. Exons were grouped by gene using the exon list for the reference GRCh37 retrieved through Ensembl’s biomart tool.

## 3 Results

### 3.1 Tests on simulations

#### 3.1.1 Model choice: detecting the presence of sex chromosomes

We used simulated data to test the range of validity of SDpop with a controlled population genetic background. We used varying numbers of individuals, different fractions of sex-linked sequences in the genome and varying times since recombination suppression, excluding biologically implausible scenarios (*i*.*e*. a very small fraction of sex-linked sequences and long times since recombination suppression, or the inverse). The results are shown in Fig. 1 and Supp. Fig. 1.

**Figure 1:**
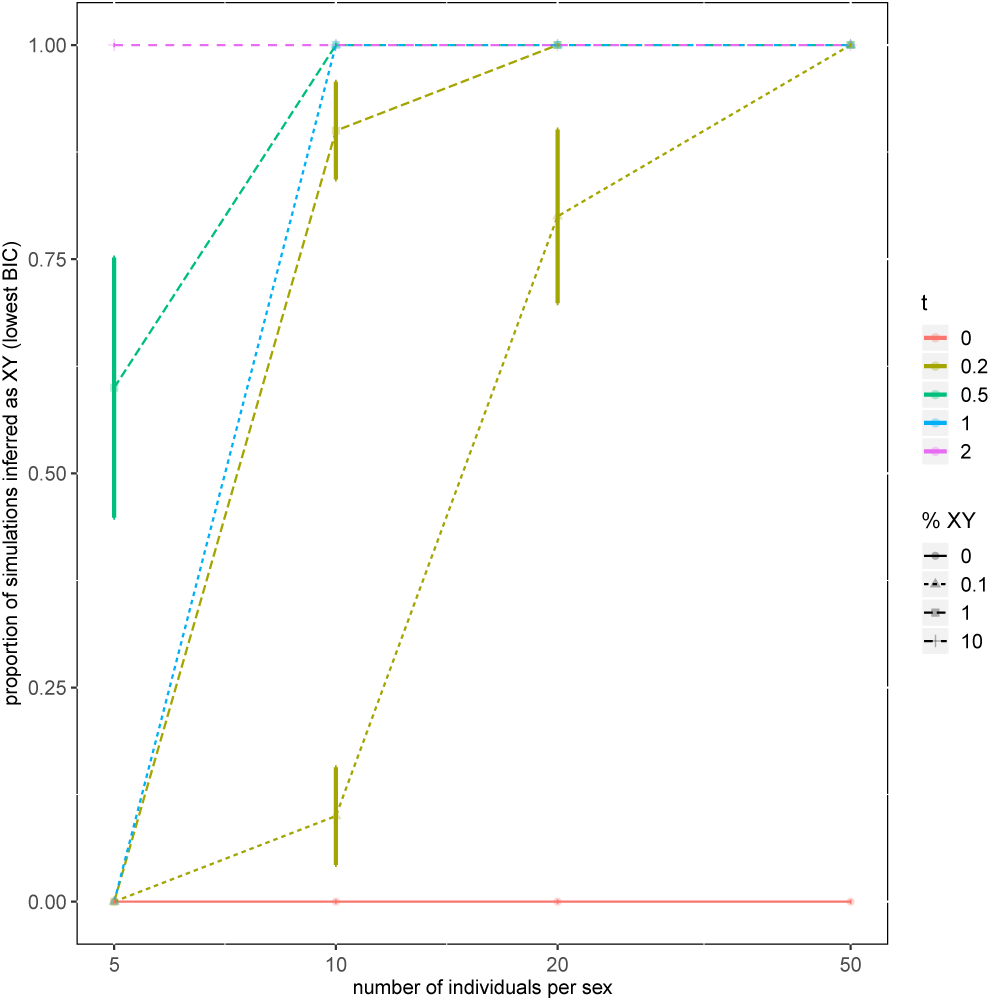
Model choice by SDpop on simulated data. The proportion of simulations for which the XY model had the lowest BIC is indicated; each combination of simulation parameter values was repeated 10 times from a random seed. Colors represent the time since recombination suppression (*t*) and the line types the percentage of genes that was simulated as gametologous. For clarity, a subset of the data, notably only with *e* = 0.0001, is shown; more results are shown in Supp. Fig. 1.

The behavior of the model when only 2 individual per sex are used is somewhat erratic, and leads to type I errors (Supp. Fig. 1), which is perhaps not surprising given the information that is present in a sample of only 4 individuals. With 5 or more individuals per sex, no type I errors were observed, and the power to detect sex-linkage increases with the proportion of sex-linked sequences and the time since recombination suppression, as expected (Fig. 1). Indeed, SDpop relies on polymorphic sites that show evidence for sex linkage: for gametologs that have stopped recombining 4*N*_*e*_ generations ago (*t* = 1; this is the average time to fixation in a neutral model), we expect and observe that most “’ true”’ polymorphisms (i.e. those without errors) in sex-linked genes are polymorphic in only one of the gametologs, or have different alleles fixed on both gametologs. However, for recombination suppression much less than 4*N*_*e*_ generations ago, most polymorphisms in gametologs are either derived from ancestral polymorphism, or due to recent mutations with low alternative allele frequency in one of the copies, making detection of sex-linkage much harder. Furthermore, the number of sex-linked genes is typically low in such situations. Our simulations indicate that even with time since recombination suppression as low as 0.1 × 4*N*_*e*_ generations and 0.1% of sex-linked sequences, the method can nevertheless select the appropriate model in most cases when 50 or more individuals per sex are used (Supp. Fig. 1).

#### 3.1.2 Assignment of genes to segregation types

Contigs are considered sex-linked when their posterior probability to be either XY or X-hemizygous exceeds the threshold value of 0.8. We measure the precision of this assignment, i.e. the fraction of contigs assigned as sex-linked that were indeed simulated as sex-linked contigs, as quantified by the positive predictive value (PPV), and the power of the assignment, i.e. the fraction of contigs that were simulated as sex-linked that we are able to detect, as quantified by the true positive rate (TPR). Results are shown in Fig. 2 and Supp. Fig. 2.

**Figure 2:**
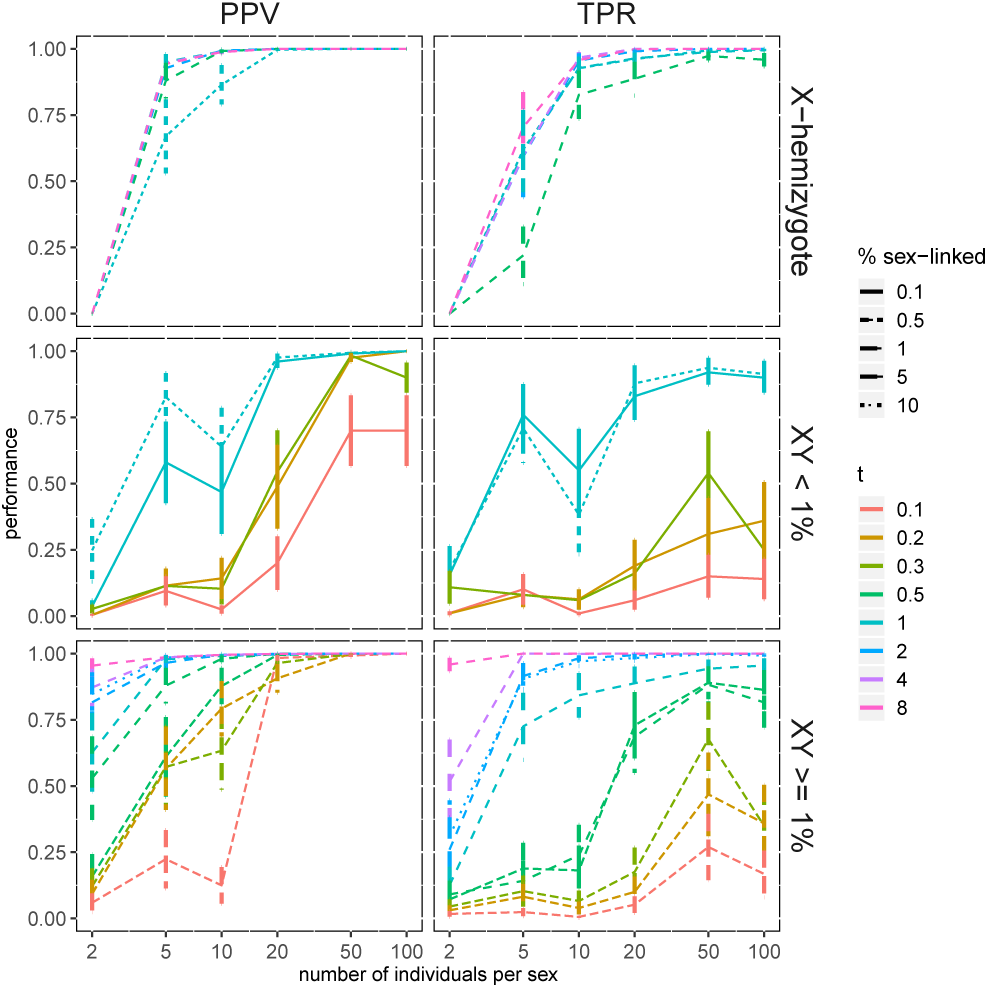
Precision (Positive Predictive Value, left) and power (True Positive Rate, right) of the detection of sex-linked contigs in simulated data, using a threshold for the posterior probability of 0.8. Top graphs: X-hemizygous genes. Middle graphs: XY gametologs, simulations with a small fraction of gametologs in the genome (< 1%); bottom graphs: XY gametologs, simulations with a larger fraction of gametologs in the genome (≥ 1%). Each point is the average of 100 simulations, with the bars representing the standard error. For all cases shown here, the simulated error rate was 0.0001; for *e* = 0.001, see Supp. Fig. 2.

For XY gametolog pairs, precision and power increase with time since recombination suppression and the size of the non-recombining region. When the time since recombination suppression exceeds 4*N*_*e*_ generations (i.e. *t* ≥ 1) and the non-recombining region is sufficiently large (i.e. more than 1% of the genome), the precision is larger than 95% and the power larger than 70% with as less as 5 individuals per sex (Fig. 2). Even with relatively recent recombination suppression (2*N*_*e*_ generations ago, i.e. *t* = 0.5), the method has a precision close to 100% and a power close to 70% with 20 individuals per sex. For shorter time since recombination suppression, both decrease rapidly. Indeed, in these cases, X- and Y-linked SNPs will still have similar frequencies, and many individuals are needed to test whether the observed allele frequency differences are due to sampling or not. For almost all most cases, the precision is higher than the power, meaning that the type I error (false positives) is lower than the type II error (false negatives).

The time since recombination suppression has no clear effect on the detection of X-hemizygous sequences, understandably, as it is only the nucleotide polymorphism in X-linked sequences that creates a signal for detection, and this level of polymorphism only depends on *N*_*e*_. Of course, when the error rate increases (Supp. Fig. 2), power and precision decrease, as expected.

#### 3.1.3 Population genetic inferences

An original feature of SDpop is that for genes inferred as gametologs, the parameter estimates and posterior probabilities can be used to estimate the allele frequencies on the X and Y copies. We use these to calculate the level of diversity in X and Y sequences, as well as their divergence. In Fig. 3, the estimates of nucleotide diversity and divergence calculated directly from the simulations, before errors were added, are compared to the estimates based on SDpop’s output.

**Figure 3:**
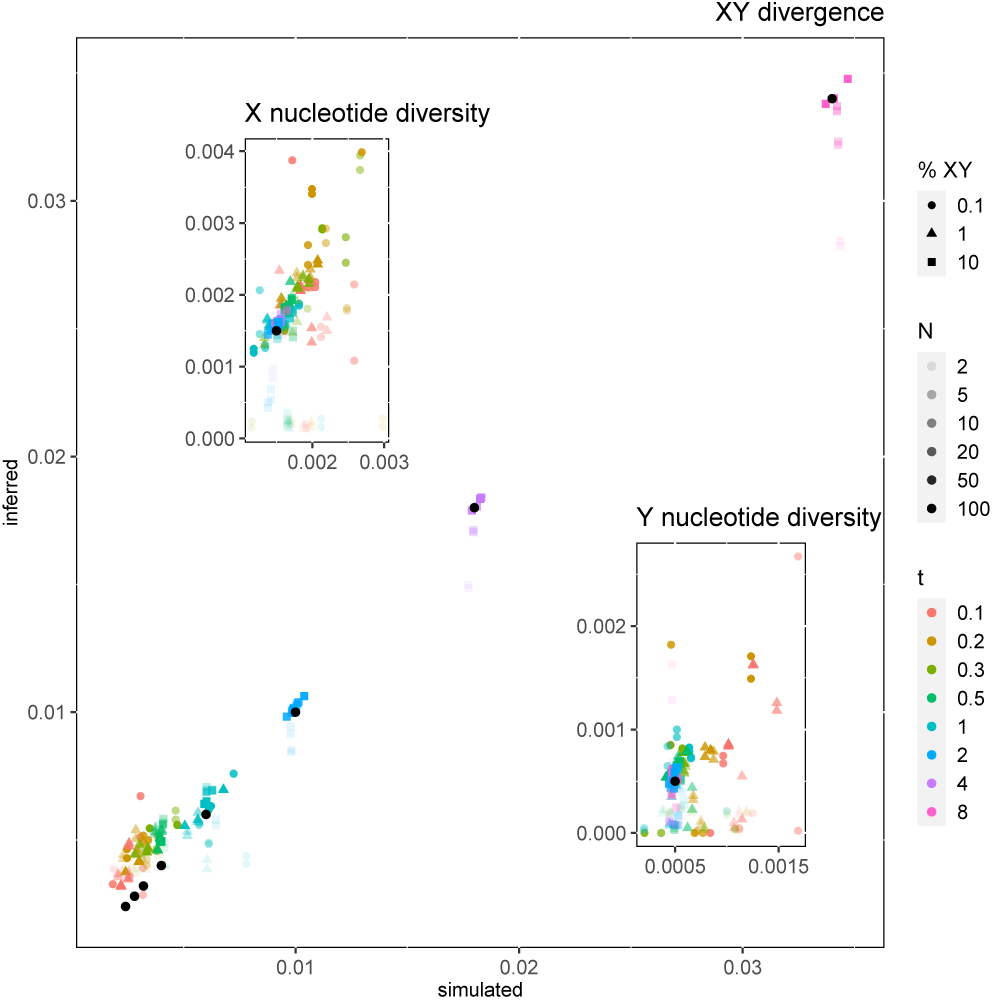
Population genetic inferences of SDpop. The values of nucleotide diversity and divergence calculated directly from the simulation results are compared to the values inferred from SDpop’s output. Comparisons are based on SDpop’s assignment of the genes (i.e. all genes with a posterior probability > 0.8 were used). Main graph: gametolog divergence (*D*_*XY*_); left inset: X nucleotide diversity (*π*_*X*_); right inset: Y nucleotide diversity (*π*_*Y*_). *N* is the number of individuals per sex, *t* the time since recombination suppression, and %XY the simulated proportion of gametologs. The black points indicate the theoretical values (*D*_*XY*_ : one for each *t*; *π*_*X*_ and *π*_*Y*_ : one value for all runs). Here, *e* = 0.0001; for higher error rates, see Supp. Fig. 3.

*D*_*XY*_ is expected to increase with time since recombination suppression, and for *t* ≥1, the values based on SDpop’s inference correlate well with the simulated values. For more recent recombination suppression, SDpop slightly overestimates *D*_*XY*_; indeed, genotyping and sequencing errors have a larger influence here. Importantly, for *t* < 1, as for *π*_*X*_ and *π*_*Y*_ the effect of random variation is large, as expected due to stochastic effects (Takahata and Nei, 1985).

### 3.2 Performance on human data

The human genome has a strongly heteromorphic XY chromosome pair, with the Y chromosome being much smaller than the X. This difference is due to the strong degeneration the Y chromosome has been subject to: it has lost many genes. Recombination suppression has occurred several times in the lineage leading to humans, leading to distinct strata characterized by different levels of degeneration (Lahn and Page, 1999). Only in the youngest stratum, genes have retained both X and Y copies, while in the older strata, Y copies have mostly been lost. Note that recombination was suppressed much longer than 4*N*_*e*_ generations ago. The most recent stratum is estimated to have stopped recombining between 30 ×10^6^ and 50 × 10^6^ years ago (Lahn and Page, 1999), while a gross estimate of 4*N*_*e*_ would be 10^6^ years (the human *N*_*e*_ has varied greatly, but is in the order of magnitude of 10^4^ (Auton et al., 2015), and the generation time is about 25 years); *t* is thus much larger than in the cases we’ve simulated.

SDpop clearly identifies the XY chromosome pair in human sequencing data (Fig. 4; Supp. Fig. 4). Here, the reference consists of the 22 chromosomes and the X (excluding the Y). The larger part of the X chromosome is detected as X-hemizygous, as most of the genes on the X have lost their Y copy. Genes on the extremities of the X chromosome have a high probability to be autosomal, which is again expected as these regions are pseudo-autosomal and do recombine between X and Y. Only one small region with XY gametologs is detected, near the left pseudo-autosomal region, at the position of the youngest stratum with XY gametologs. The genes in the older strata, for which the Y copies have been lost, are detected as X-hemizygous by SDpop. The Y also has several genes without X homologs, that probably resulted from transpositions or translocations from other autosomes, the so-called ampliconic genes (Skaletsky et al., 2003). Indeed, the autosomal gene DAZL, known to have given rise to the Y-ampliconic family DAZ, had a very high probability to be sex-linked (> 0.99). Other genes are also shown be homologous with sequences on the sex chromosomes (Galichon et al., 2012), and two of these (PPP1R12B and TPTE2) harbor the majority of sex-linked SNPs detected outside chromosome X.

**Figure 4:**
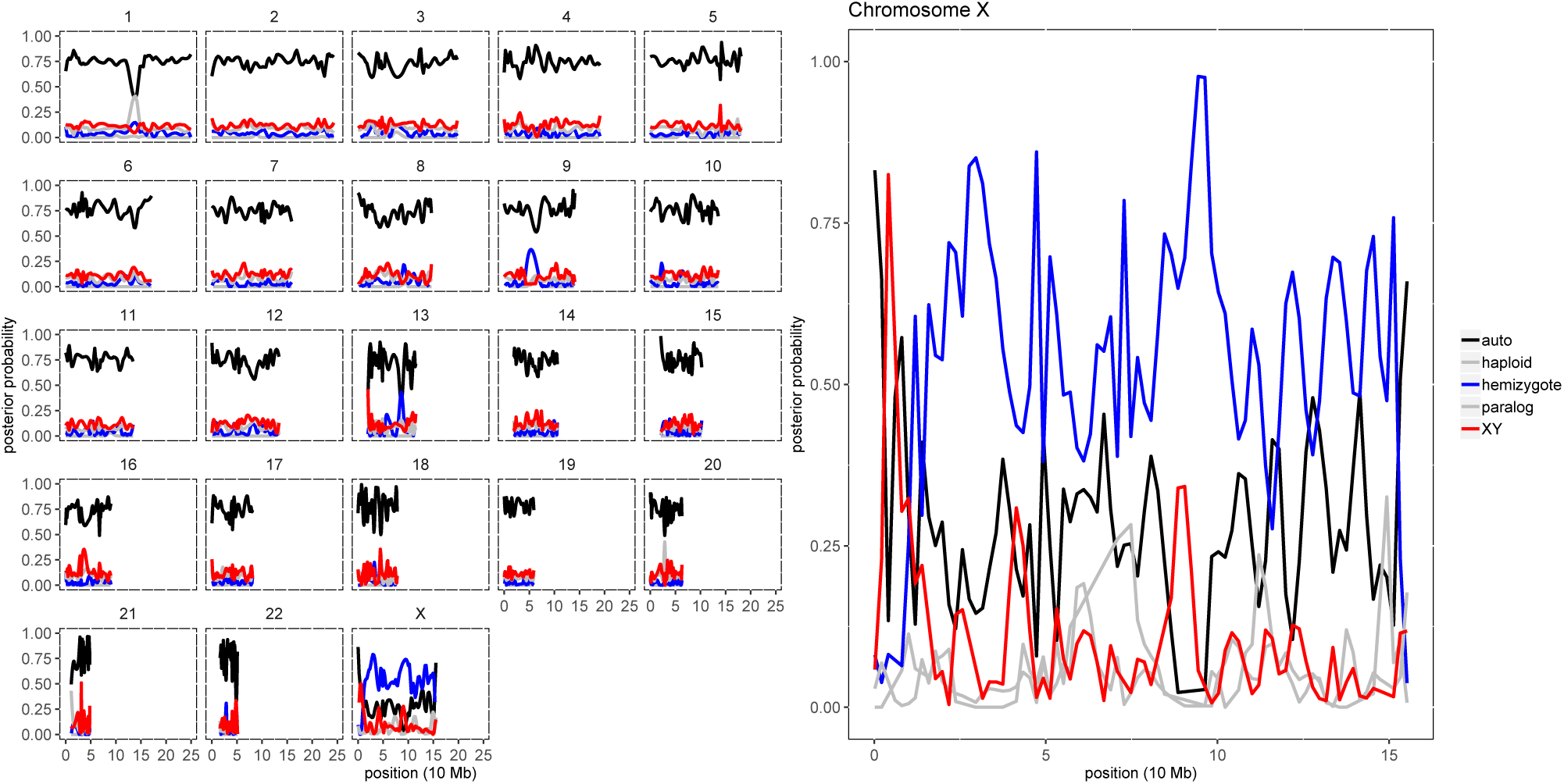
Test of SDpop’s performance on the human exome-targeted sequencing data from the 1000 genome project, using 10 individuals per sex. Smoothed gene-level posterior probabilities for autosomal (black), X-hemizygote (blue) and XY (red) segregation are shown; haploid and paralogous posterior probabilities are indicated in grey. The right panel show the results on the X chromosome: the extremities corresponding to the pseudo-autosomal regions are predicted to be autosomal by SDpop, while most XY gametologous genes are found close to the pseudoautosomal region on the left arm, which represents the youngest stratum where Y copies have not yet been lost. The rest of the chromosome consists of X-hemizygous genes, that lack a Y copy. Results obtained with 5 and 20 individuals per sex are shown in Supp. Fig. 4.

As Y-specific genes are often present in a few copies, which allows gene conversion to rescue these sequences that suffer from the lack of recombination, Y-specific diversity and XY-divergence are not correctly estimated from the outputs of SDpop. Indeed, we observe that the parameter *ρ*, the proportion of X-polymorphism in XY gametologs, is 0.12 when using 20 individuals per sex, indicating that 88% of polymorphism in XY gametologs is due to Y polymorphism. This could be due to the mapping of several copies of the Y gene to the same position on the X chromosome.

## 4 Discussion

SDpop is a probabilistic framework for the detection of sex-linked sequences, that relies on the modeling of the expected equilibrium between allele and genotype frequencies under sex-linked segregation. It combines the principles that are at the basis of several methods for detection of sex-linkage, such as increased frequencies of heterozygotes or allele frequency deviations, in a specific framework. As such, it requires less individuals than methods that are based on such allele frequencies or genotype frequencies alone: e.g., GWAS usually needs > 50 individuals, as do studies using tests for heterozygote frequencies (Picq et al., 2014). Of course, SDpop’s power depends on the size and age of the non-recombining region, as would be the case for any method.

The likelihood-based framework of SDpop allows to compare models with and without sex linkage. It is thus possible, with a moderate sequencing effort, to determine the sex chromosome system and to obtain the sex-linked sequences for any species that can be sexed and sampled in the field. The approach can be used on any kind of individual-based sequencing data, such as RNA-seq, DNA-reseq and RAD-seq. The functionality to calculate gene-level posterior probabilities is of course only useful for gene-based sequencing, such as RNA-seq and exome capture. For DNA-reseq, posterior probabilities could be calculated when the scaffold size is small, or by splitting the chromosomes into windows of fixed size.

The underlying population genetics model uses the classical assumptions of the Hardy-Weinberg principle, notably random mating between the sexes in an infinitely large population. The model proposed here will thus perform best when used on a sample of individuals taken from a single, panmictic population. Any population structure will weaken the performance of the model. Indeed, when females and males are sampled from separate populations, this will lead to type I errors (false positives), as the population structure mimics the deviations from Hardy-Weinberg equilibrium that are expected for sex linkage. However, if the individuals sampled from different populations comprise both females and males from each population, the expected influence on SDpop’s performance is mainly a loss of power (i.e., an increased proportion of false negatives).

We model different kinds of sources of error in SDpop. First, we explicitly model haploid and paralogous sequences. These are more or less frequently encountered in NGS data; haploid sequences could result from contamination (e.g. mitochondria, chloroplasts, or bacteria), monoallelic expression in RNAseq data, or redundancy in the reference (i.e. different alleles of a gene are split into several contigs). Paralogous sequences could result from recent paralogs that were not recognized as such in the reference genome or transcriptome, or from contamination between samples. Furthermore, SDpop incorporates an error parameter, to account for other sequencing or genotyping errors. These sources of errors are modeled solely to improve the detection of sex-linkage: failing to take them into account would lead to haploid sequences being inferred as X-hemizygotes, paralog sequences as XY gametologs, while sequencing and genotyping errors would penalize these sex-linked segregation types more than the inference of autosomal segregation. The power, sensitivity and precision of SDpop’s inferences of haploid and paralogous segregation as well as the error rate have not been tested extensively, and thus one should not use the method to estimate these segregation types and the error rate from the data.

SDpop’s sex-linked segregation types include both gametologous segregation and hemizygote segregation. In the first case, both gametologous copies are present, while in the second, there is no information about the sequence from the sex-limited chromosome. X-hemizygous loci thus correspond loss of the Y copy of a gene, presumably through Y degeneration. However, apparent X-hemizygosity can also be caused by artifacts that are more or less difficult to control. First, the X and Y copies might be incorporated as distinct genes in the genome or transcriptome assembly. In species with known and well described sex chromosomes, such as humans, the Y assembly could simply be excluded from mapping, as we did here. In species with more recently evolved or less well described sex chromosomes, one should thus preferably use the homogametic sex for preparing a mapping reference. The case that is harder to solve arises when Y sequences are too divergent to map on the X copy in the reference; thus, when some sex-linked genes have high XY divergence values, one should be aware of the fact that some genes that have been inferred as X-hemizygous might be artifacts.

As shown here and by others (e.g. Crowson et al., 2017), XY gametologs are much easier to detect than X-hemizygous genes, as, first, XY gametologs will have more SNPs than X-hemizygous genes, and second, the information contained in a fixed XY SNP is much less ambiguous than the one of a X-hemizygous SNP. These are additional reasons to try to reduce the number of artifactual X-hemizygous as much as possible, in order to obtain more reliable inferences.

Importantly, there is further information to be obtained from XY gametologs. We use the inferred allele frequencies on the X and the Y copies to calculate the nucleotide diversity in the X and Y copies, and their divergence. Thus, using the information on which alleles are fixed in each copy, SDpop is also able to reconstruct the haplotypes of the X and Y copies, even when the input data have not previously been phased. We’ve shown that the estimation of population genetic parameters comes quite close to the simulated values, even though these estimations are based on empirical allele frequencies (i.e., they do not take the error rate into account). However, even if the estimate would be perfectly unbiased, we expect that the variance of these parameters on a per-gene basis (cf Takahata and Nei, 1985) is much larger than the bias introduced by not taking the error rate into account. Thus, obtaining reliable estimates of these parameters (i.e., estimates that reflect population processes and not mere stochasticity) requires many independent samples, and cannot be addressed solely by modeling.

Although it has similarities with available methods, SDpop is unique in the combination of input data it requires and the predictions it can produce. As a consequence, we cannot compare its performance directly to any of previously published methods. Our simulations and tests show that reasonable performance can be achieved with as few as 5 individuals per sex. This is close to the sequencing effort required for SEX-DETector (Muyle et al., 2016), which relies on a controlled cross to infer sex-linkage. Given the fact that usage of SDpop puts less constraint on the input data (kinship does not need to be known, and is assumed to be absent), it is, at the first sight, somewhat surprising that SDpop can work with as few individuals as can SEX-DETector. The reason for this is that not all sites are informative for SEX-DETector (e.g. when both parents are heterozygous) and are discarded, whereas SDpop considers all polymorphic sites as informative; the amount of information supplied by a site varies depending on the difference of the conditional likelihood for all segregation sites.

Apart form the main practical advantage that SDpop does not require controlled breeding of the study organism, it has a few more advantages compared to SEX-DETector. First, as SEX-DETector considers parents and their F1 offspring, there has been little recombination between the homologous chromosomes. For the sex chromosomes, this implies that genes from the pseudo-autosomal region are genetically linked to the non-recombining region, and will have more or less distorted segregation. This leads SEX-DETector to overestimate the size of the non-recombining region, especially when the pseudo-autosomal region is large and the sex-linked region small (with a large non-recombining and a small pseudo-autosomal region, this effect is less important, as the sparse recombination events of the sex chromosomes will be located in the small pseudo-autosomal region, and linkage disequilibrium will decrease rapidly). Thus, we expect SDpop to yield a more precise indication of the pseudo-autosomal boundary. Note that, in the case of a very small and recently evolved non-recombining region that will be difficult to detect by SDpop, SEX-DETector’s behavior of overestimating the non-recombining region might be beneficial as it increases the capacity to detect the sex chromosome; in such cases, model choice could be done with SEX-DETector on a controlled cross, and delimitation of the non-recombining region with SDpop on population data.

Second, an advantage of the use of population data in SDpop is that estimates of population genetic parameters are possible. In a cross, there will be three X chromosomes and one Y, so it will be impossible to distinguish fixed substitutions from polymorphism on the Y chromosome.

DETSEX (Gautier, 2014) uses a Bayesian framework that can be used on samples collected in natural populations to infer whether markers (SNPs) are sex-linked or not. Despite these obvious similarities with the model proposed here, there are several important differences. First, SDpop allows to compare of the total likelihoods of the models, and thus can be used as a statistical test for the presence or absence of sex chromosomes, while in DETSEX, the presence of sex chromosomes is assumed, but the individual’s sex does not need to be known. Second, in DETSEX, X and Y-linked sequences are expected to map to different positions, which can be a safe assumption in well-studied species with an old sex chromosomes system (such as humans), but not for more recently evolved sex chromosomes.

Thus, SDpop fills a gap in the current panel of methodological approaches for the identification and study of sex chromosomes. It requires input data that have now become classical: short reads sequencing of genomes or transcriptomes of ten to twenty individuals, collected in any population. It uses standard vcf files as input, thus allowing integration in existing genotyping pipelines. Its probabilistic framework and its implementation in a widely used and efficient programming language (C/C++) furthermore allow future developments (including, but not limited to, inference of individual’s sex, corrections for population structure).

## Acknowledgements

We thank Laurent Duret for help with the treatment of the 1000 genomes data, François Gindraud for help with programming, and Sylvain Mousset and Thibault Latrille for discussions on population genetics issues. This work was performed using the computing facilities of the CC LBBE/PRABI, and was supported by funding from the Agence Nationale de la Recherche (grant number ANR-14-CE19-0021-01).

## Author contributions

GM conceived the project and acquired funding; JK, NL, GM and FP conceived the model; JK, NL and FP developed the methodology and the formal analysis; JK developed the software, performed simulations and data analysis, and prepared the first draft manuscript; JK, GM and FP wrote the current version of the manuscript; all authors approve of the manuscript.

## Appendix

### Full description of the model

#### Observed and hidden variables

The data consist of genotyped individuals for genes *k*∈ {1, 2, …, *K*}, each with biallelic sites *t* ∈{1, 2, …, *T*_*k*_}. For each biallelic site, the observed alleles are randomly named *a* and *b*, such that three genotypes are possible: homozygote for allele *a* (“*aa*”), heterozygote (“*ab*”), and homozygote for allele *b* (“*bb*”). At each biallelic site *t* of gene *k*, and for each individual *i*, 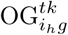 is an indicator of the individual having observed genotype *g* ∈ {1 = *aa*, 2 = *ab*, 3 = *bb*} and sex *h*∈ {1, 2}. *h* = 1 for females, *h* = 2 males; for convenience, we will write ♀ and ♂. 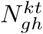 is the number of individuals with sex *h* and observed genotype *g*; note that the total number of observations (i.e., genotyped individuals) can vary between sites. The vector **OG**^*kt*^ describes all observations at a site.

We seek under which segregation type *S*_*j*_, *j* ∈ 1…7, these observed genotypes are most likely. These segregation types will be listed in fixed order, and a specific number corresponds to each of them:

1. Diploid autosomal segregation
2. Haploid sequences
3. Paralogs
4. X-hemizygous segregation
5. XY gametologous segregation
6. Z-hemizygous segregation
7. ZW gametologous segregation

For some segregation types, as will be detailed below, several sub-types of segregation have to be specified; these are denoted *A*_*l*_ with *l* ∈{1..*L*}. The conditional likelihood of observing the genotypes under each segregation type depends on the allele frequencies and a genotyping error rate.

We introduce a hidden variable 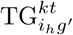 which is an indicator for the true genotype *g*′ ∈{*aa, ab, bb*} of an individual *i* with sex *h*. The conditional probabilities of observing a true genotype for an individual, given the fully specified segregation type, are

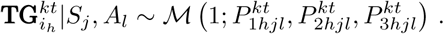

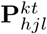 is the vector of the probabilities 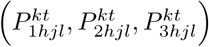 for each genotype at a site, given the sex of the individual and the segregation type and subtype. The genotype probabilities 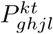 are calculated from the empirical allele frequencies 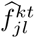 using the following population genetic expectations.

1. For autosomal segregation, the Hardy-Weinberg equilibrium should hold in both sexes. Thus,

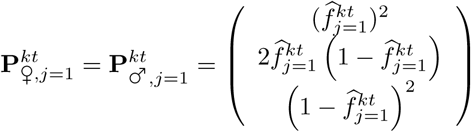

where

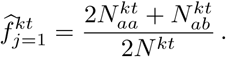
2. Haploid segregation is modeled by

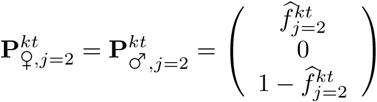

and

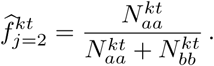
3. Paralogy is caused by the mapping of the reads of two more or less recently duplicated genes on one locus in the reference. There is no recombination between the copies, that thus evolve independently. For simplicity, we assume that one of the copies is fixed for one of the alleles. The genotype probabilities depend on which allele is considered fixed in one of the copies, and two sub-types have to be modeled.
  a. First, we consider that allele *a* is fixed in one of the copies. 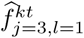 is the frequency of allele *b* in the other copy. In reality, such sites have four copies; thus, the genotypes are *aaaa, aaab*, are *aabb*, with frequencies 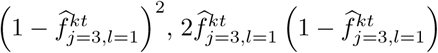 and 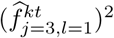. Genotypes *aaab* and *aabb* will probably be considered as *ab* by the genotyper that expects only diploids; thus, the genotype probabilities are:

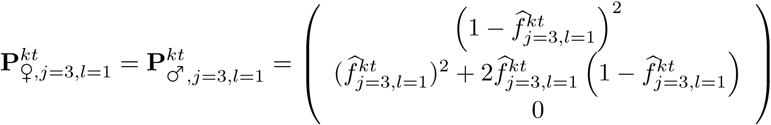 To estimate the empirical allele frequency, note that the *ab* genotype counts that are obtained from the genotyper results will be a mixture of *aaab* and *aabb*, in the expected proportions that depend on the frequency that we concisely denote 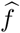 here: 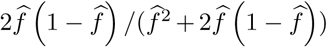 and 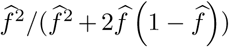. While in reality, 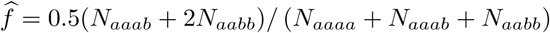, we instead calculate

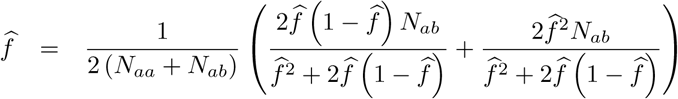

This yields

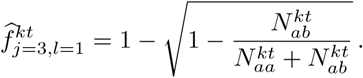
  b. Alternatively, allele *b* could be fixed in one of the copies. 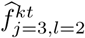 is the frequency of allele *a* in the other copy. The genotype probabilities and empirical allele frequency are

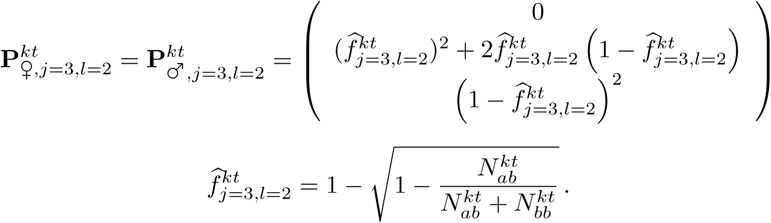
4. For X-hemizygously segregating genes, the members of one sex are haploid while the others are diploid.

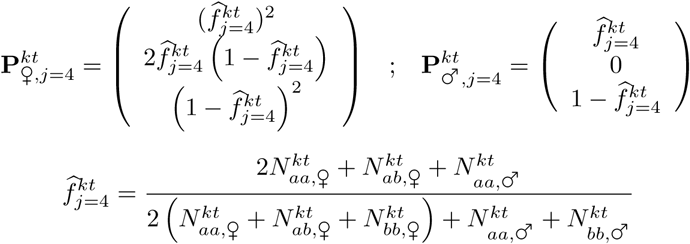
5. XY gametologous segregation is characterized by the presence of two independent copies in males, and two copies of the X gene in females. We assume that an allele is fixed in at least one of the copies.
  a. X-polymorphism, allele 1 fixed on Y. *f* is the frequency of allele 2 on X.

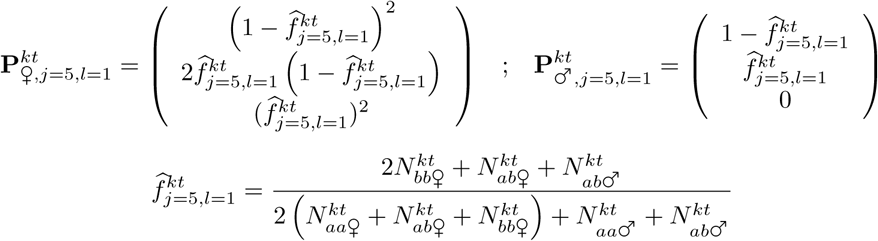
  b. X-polymorphism; allele 2 fixed on Y. *f* is the frequency of allele 1 on X :

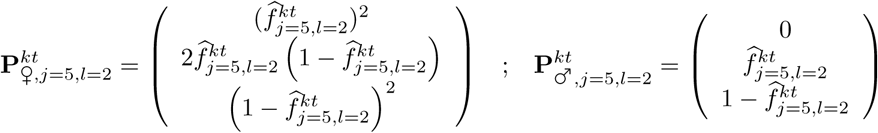

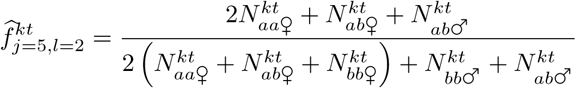
  c. Y-polymorphism, allele 1 fixed on X. *f* is the frequency of allele 2 on Y:

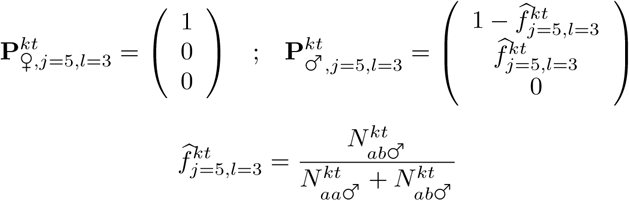
  d. Y-polymorphism, allele 2 fixed on X. *f* is the frequency of allele 1 on Y:

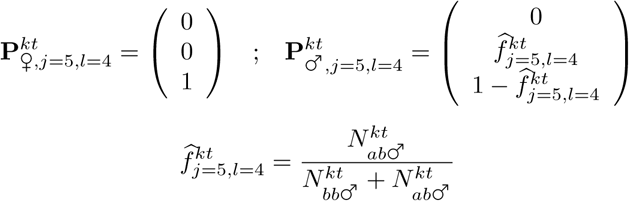
6. Z-hemizygous segregation is similar to X-hemizygous segregation:

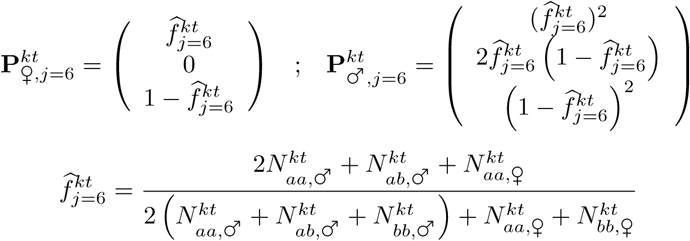
7. ZW gametologous segregation is modeled similar to XY gametologous segregation, for both Z and W polymorphism, and two asymmetrical cases for each.
  a. Z-polymorphism, allele 1 fixed on W. *f* is the frequency of allele 2 on Z.

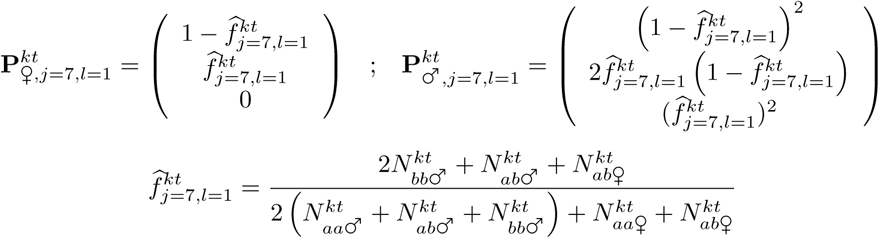
  b. Z-polymorphism; allele 2 fixed on W. *f* is the frequency of allele 1 on Z:

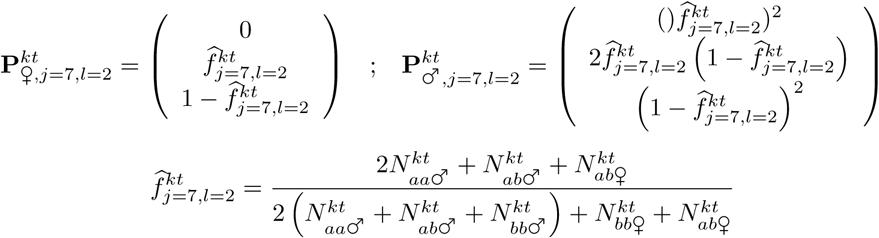
  c. W-polymorphism, allele 1 fixed on Z. *f* is the frequency of allele 2 on W:

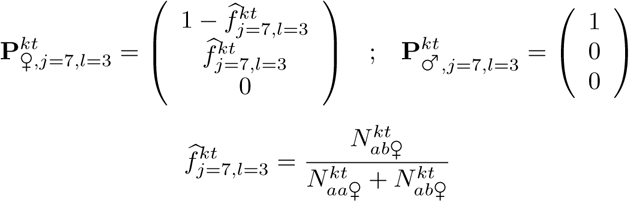
  d. W-polymorphism, allele 2 fixed on Z. *f* is the frequency of allele 1 on W:

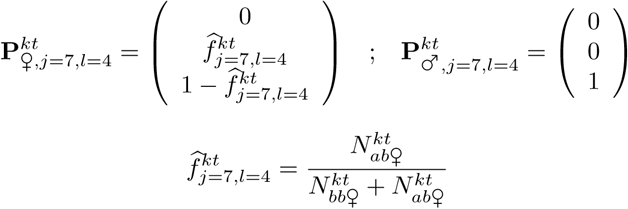

In some cases, calculation of 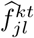 might lead to division by 0. To avoid this problem, counts that are expected to be 0 under a segregation type are added to the numerator and the denominator.

Genotyping errors (whether they are due to sequencing errors, read mapping errors, or violations of the assumptions of the method for genotyping) cause the observed genotype *g* to be different from the true genotype *g*′. We define 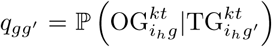, *i*.*e*., the probability to observe genotype *g* when the true genotype is *g*′, and **Q** is the matrix of all *q*_*gg*′_, such that

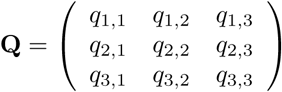

We can now directly calculate the probabilities of the observed genotypes for each segregation type:

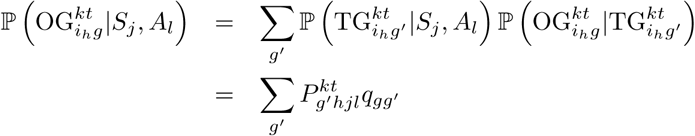

We rename the quantity 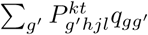 as 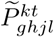; it is the expected frequency of the observed genotype given the segregation type and a certain genotyping error rate. For each sex, **OG** follows a multinomial distribution 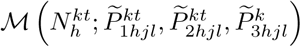. Thus, the conditional likelihood of the data (given the segregation type) at each site, that we name 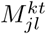, is

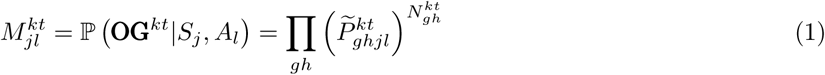

##### Parameters

The error rates *q*_*gg*′_ depend on one error parameter *e*. We assume all genotyping errors to occur with the same frequency, so *q*_*g,g*′≠ *g*_ = *e* and *q*_*g,g*′≠ *g*_ = 1 − 2*e*, which gives the error matrix

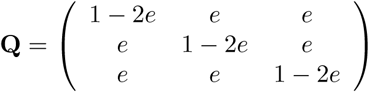

Two more series of parameters are required to model the data; these indicate the proportion of the genome that segregates under each type. There are a maximum of seven segregation types *S*_*j*_, each occupying a proportion *π* _*j*_ of the genome, such that ∑_*j*_ *π*_*j*_ *= 1*. ***π*** is the vector containing all *π*_*j*_. The segregation types 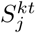 are distributed multinomially, thus

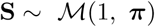

Several biologically relevant segregation types (*S*) have several “subtypes”(*A*), depending on the fixation of one of the alleles on either of the copies. Thus, for a segregation type *S*_*j*_, there are *L* subtypes, and each subtype *A*_*jl*_ applies to a proportion *α*_*jl*_ of the proportion *π*_*j*_ of the genome (corresponding to the segregation type *S*_*j*_). For each segregation type with subtypes, 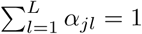, and

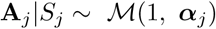

For the paralogs, the subtype depends uniquely on the choice of what allele is called *a*, which is random. Thus, no parameter is needed, and

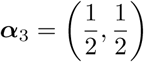

For the XY and ZW types, more sites can be polymorphic on one chromosome than on the other. The proportion of XY or ZW sites that are polymorphic on X or on Z is described by the parameter *ρ*_*j*_, which takes a single value for each segregation type. The (random) choice what allele is called *a* affects both X (or Z) and Y (or W) polymorphisms, leading to four subtypes

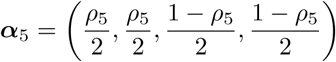

When aggregating the segregation subtypes (*A*) to biologically relevant types (*S*), we get

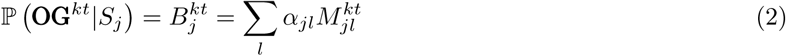

##### Expectation-Maximization algorithm

The full log-likelihood of the model is given by

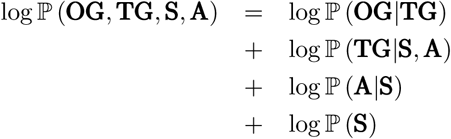

This likelihood is maximized through an Expectation-Maximization (EM) algorithm.

###### E-step

The posterior segregation types are given by

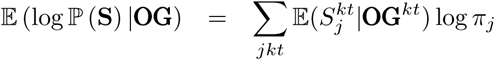

with

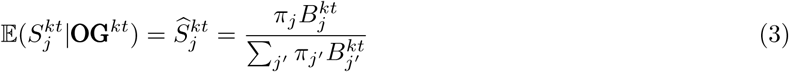

The posteriors for the subtypes are calculated by

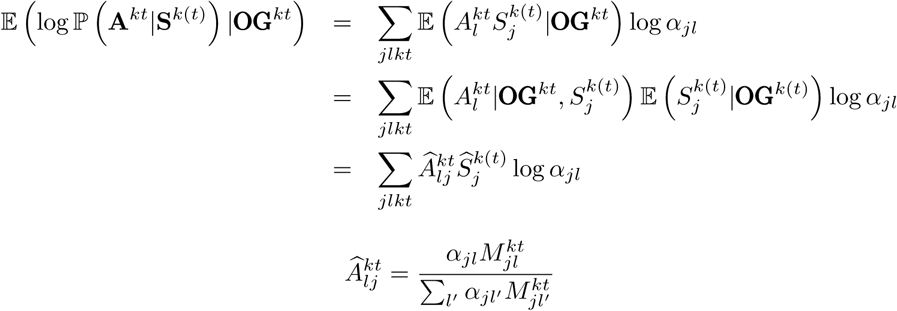

For the true expected true genotypes, we calculate

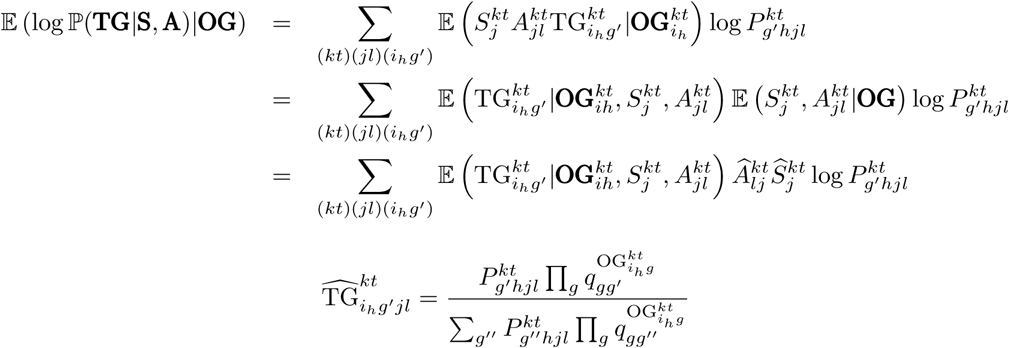

As individuals are defined uniquely by their sex and their observed genotype,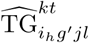 is the same for two individuals having the same sex and genotype. Thus, we write 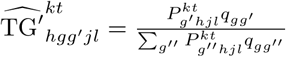.

Finally, the conditional likelihood of the observed genotypes is given by

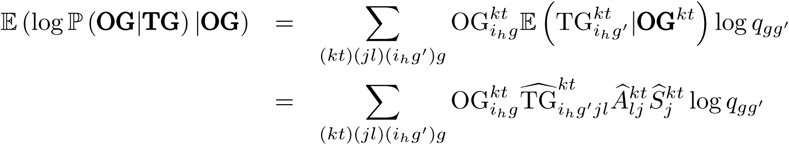

###### M-step

The key quantity to be used in the M-step is the conditional expectation of the complete-data likelihood:

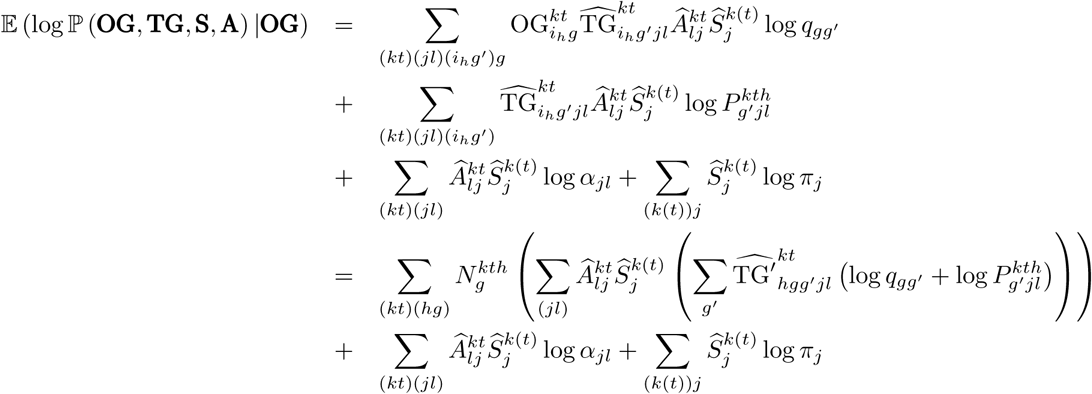

Parameters to estimate are ***π, α*** and error rate *e*. These parameters only involve

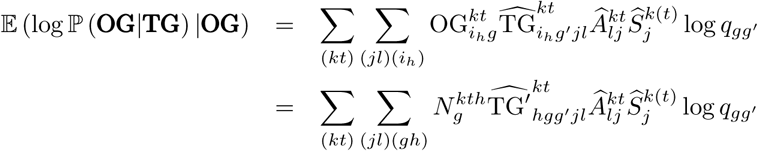

To simplify notations, let us denote

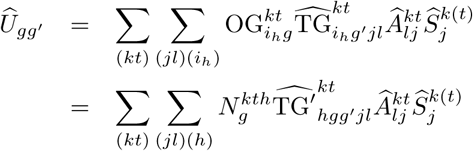

Thus,

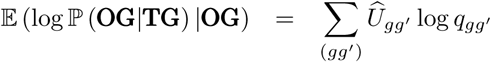

We find the new values of *e* by 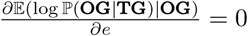, which gives:

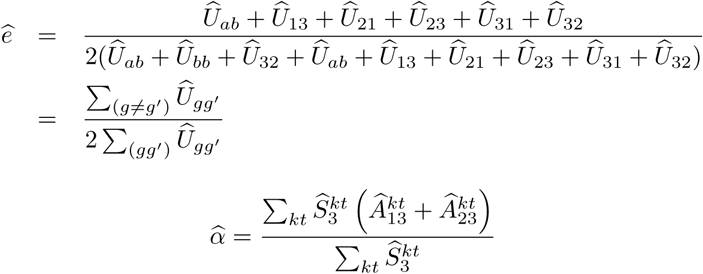

###### Monitoring and convergence

The likelihood of the data in the model is

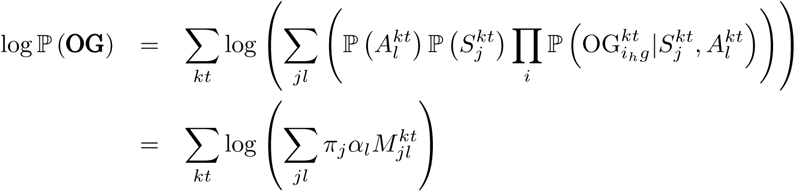

Convergence is evaluated as a function of the relative change in parameter value estimations. Optimization is halted when the largest relative change of all parameters has been less than 10^−4^ for 10 iterations, except for the error rate parameter, which is not considered for convergence.

There are *J* − 1 free parameters for the segregation types, one parameter *α* for each of the XY and ZW types, and one parameter for the error rate. If the number of parameters is *ν*, we calculate the Bayesian Information Criterion (BIC) as follows:

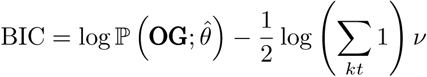

The model with the lowest BIC has the best fit.

###### Site- and contig-wise probabilities

The posterior probabilities per site, as given in Eq. [3], are calculated using the priors *π*_*j*_. The smaller *π*_*j*_, the higher the conditional likelihood 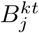 should be to produce a high posterior probability. For the sex-linked segregation types *π*_*j*_ can easily be very small. If, say, 0.1% of the sites are inferred as gametologous and 99.9% as autosomal, the conditional likelihood for the gametologous segregation types should be 1000 times higher than the one for autosomal segregation to obtain comparable posterior likelihoods with this formula. In order to avoid excessive biases against rare segregation types, for inference purposes at the end of the optimization, we calculate the posterior probabilities without priors, which amount to using a uniform prior,. Thus, for the output, we compute

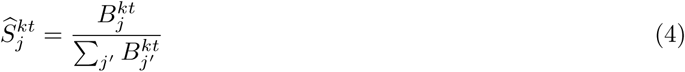

At the contig level, the goal is to estimate the posterior probability to be sex-linked, autosomal, or not informative (*i*.*e*., haploid or paralogous), given the observed data for each of its sites and the optimal parameter values. This probability is the expectation of each segregation type, 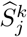. It can be calculated directly given the expectations per site, 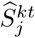 from Eq. [3]:

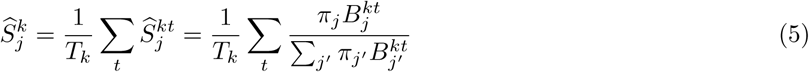

For robustness, geometric mean calculations of the posterior probabilities per contig are also calculated. The geometric mean of a series of observations 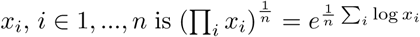. The geometric mean is more robust than the arithmetic mean since the relative value of the contig-level geometric means is independent of site-level normalizations, which is not true for arithmetic means. Furthermore, the ratio of the geometric means of two series is equal to the geometric mean of the ratios of the elements of the series. The posterior probability based on the geometric mean is

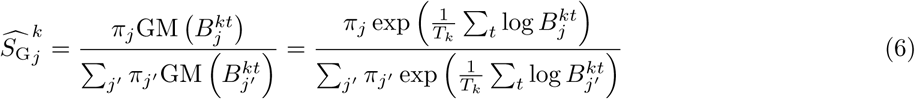

Finally, we calculate the geometric mean of the site-wise conditional likelihoods without taking the proportion *π*_*j*_ for each segregation type as a prior. Indeed, when the sex-linked region is very small, containing few fixed differences, these proportions will be very low for the sex-linked segregation types, and the influence of the prior would be too strong. Thus, for the output, we calculate:

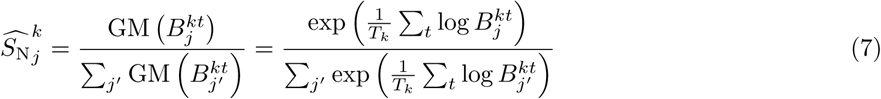

We recommend that inferences of segregation types should be based on the posterior probabilities that were calculated without the priors, *i*.*e*. Eq. [4] for sites and Eq. [7] for contigs. Other posteriors are nevertheless provided for expert users.

###### Population genetic predictions

From the allele frequencies and segregation subtypes, it is possible to calculate the expected diversity and divergence of the gametologous copies. For each site, the frequency of allele *a* on chromosome *v* ∈ {*W, X, Y, Z*} is

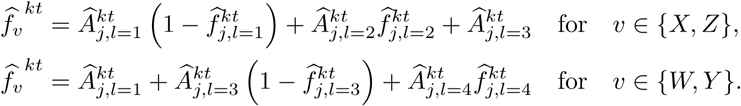

A different way of predicting the allele frequency on both sex chromosomes is to assign it to be the frequency corresponding to the most probable subtype.

This information can be used to infer the consensus sequences of the X and Y sequences. For a given contig (that can be chosen on the basis of 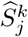, but not necessarily if we have other reasons to believe the contig is sex-linked), each polymorphic site can be considered fixed for an allele if 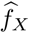 or 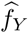 are above a threshold *U*_*f*_ (0.5 ≤ *U*_*f*_ ≤ 1) or below 1 − *U*_*f*_. A further threshold can be applied to genotype non-fixed sites: if 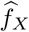 or 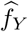 are above a threshold *u*_*f*_ (0.5 ≤ *u* _*f*_ ≤ *U* _*f*_) or below 1 − *u* _*f*_.

Nucleotide diversity can be calculated as

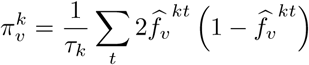

where *τ*_*k*_ ≥ *T*_*k*_ is the total length of the contig *k*, including monoallelic sites. The divergence is

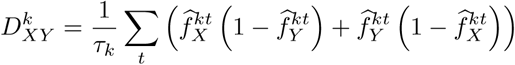

in the XY case; extension to ZW chromosomes is trivial.

## Supplementary figures

**Supplementary Figure 1:**
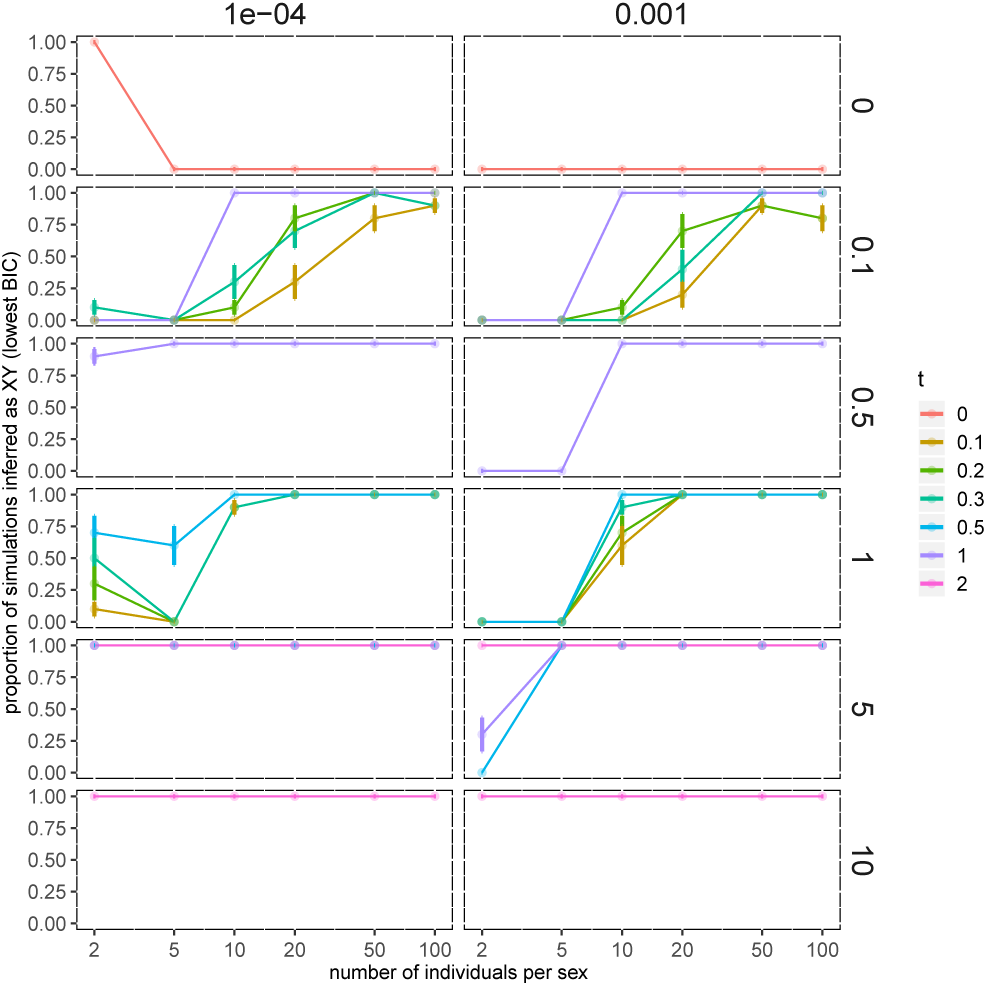
Model choice by SDpop on simulated data. The proportion of simulations for which the XY model had the lowest BIC is indicated; each combination of simulation parameter values was repeated 10 times from a random seed. Left column: simulated error rate *e* = 0.0001, right column: *e* = 0.001; the percentage sex-linked sequences varies from none (top row) to 10% (bottom).

**Supplementary Figure 2:**
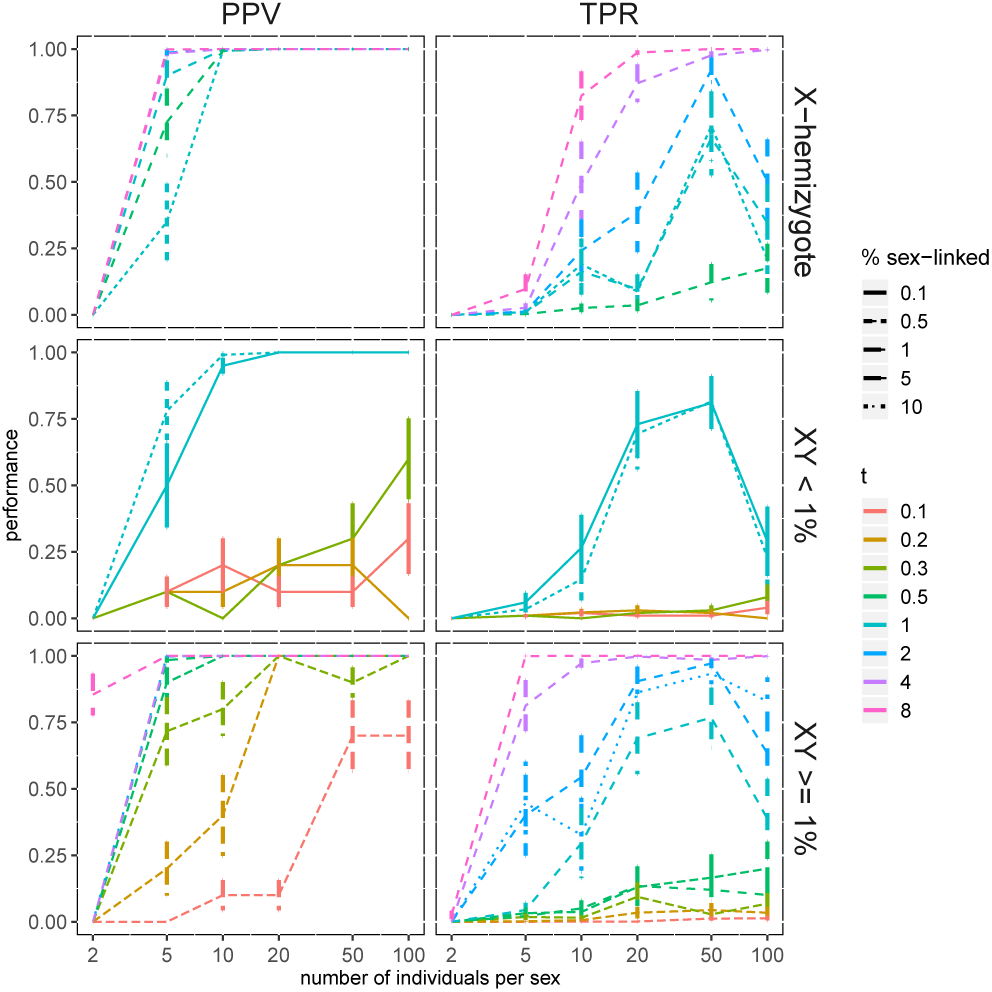
Precision (Positive Predictive Value, left) and Power (True Positive Rate, right) of the detection of sex-linked contigs in simulated data, using a threshold for the posterior probability of 0.8.. Top graphs: X-hemizygous genes. Middle graphs: XY gametologs, simulations with a small fraction of gametologs in the genome (< 1%); bottom graphs: XY gametologs, simulations with a larger fraction of gametologs in the genome (≥ 1%). Each point is the average of 100 simulations, with the bars representing the standard error. For all cases shown here, the simulated error rate was 0.001; for *e* = 0.0001, see Fig. 2.

**Supplementary Figure 3:**
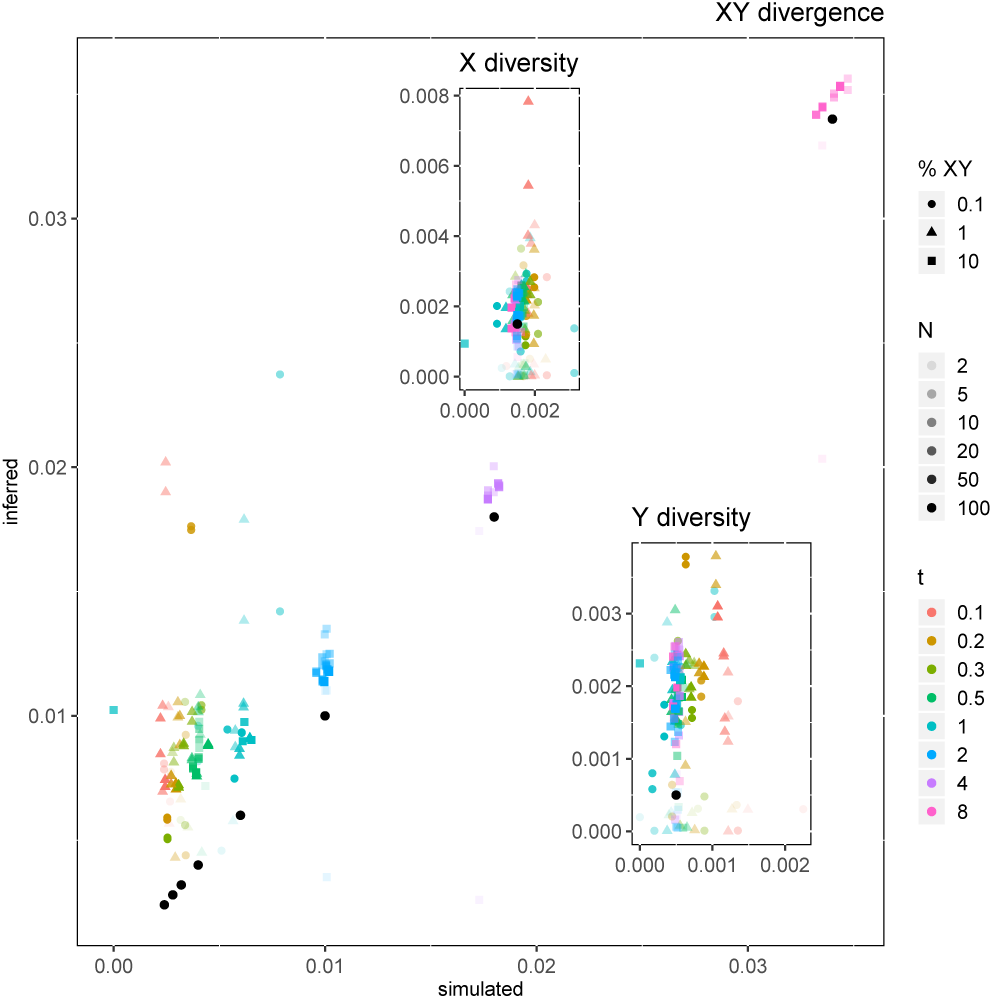
Population genetic inferences of SDpop at higher error rates than Fig. 3; here, *e* = 0.001. The values of nucleotide diversity and divergence calculated directly from the simulation results are compared to the values inferred from SDpop’s output. Comparisons are based on SDpop’s assignment of the genes (i.e. all genes with a posterior probability > 0.8 were used). Main graph: gametolog divergence (*D*_*XY*_); left inset: X nucleotide diversity (*π*_*X*_); right inset: Y nucleotide diversity (*π*_*Y*_). *N* is the number of individuals per sex, *t* the time since recombination suppression, and %XY the simulated proportion of gametologs. The black points indicate the theoretical values (*D*_*XY*_: one for each *t*; *π*_*X*_ and *π*_*Y*_: one value for all runs).

**Supplementary Figure 4:**
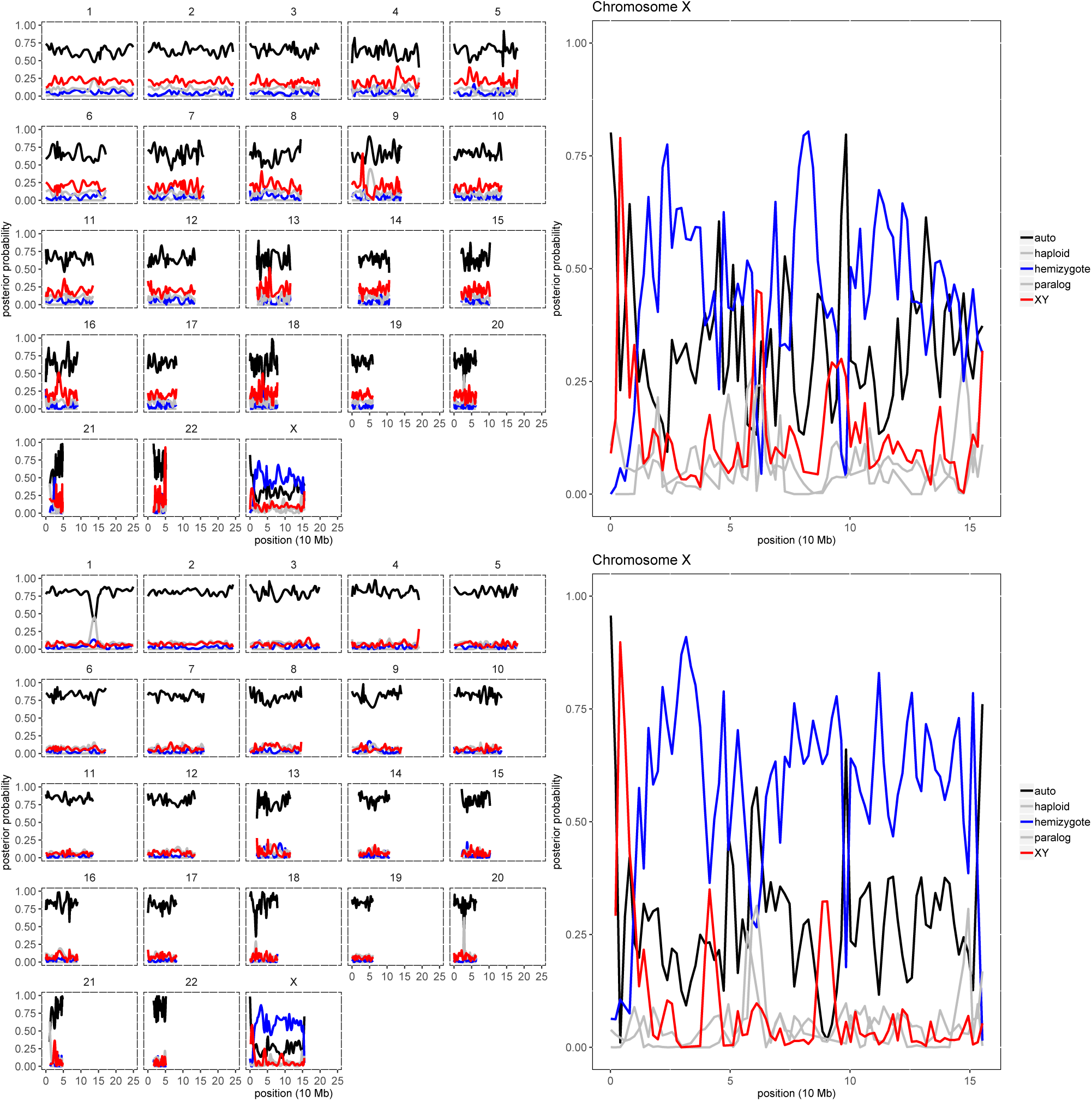
Test of SDpop’s performance on the human exome-targeted sequencing data from the 1000 genome project, using 5 individuals per sex (top graphs) or 20 (bottom graphs). Smoothed gene-level posterior probabilities for autosomal (black), X-hemizygote (blue) and XY (red) segregation are shown; haploid and paralogous posterior probabilities are indicated in gray. The right panels show the results on the X chromosome: the extremities corresponding to the pseudo-autosomal regions are predicted to be autosomal by SDpop, while most XY gametologous genes are found close to the pseudo-autosomal region on the left arm, which represents the youngest stratum where Y copies have not yet been lost. The rest of the chromosome consists of X-hemizygous genes, that lack a Y copy.

